# Collaborative multi-agent intelligence uncovers subtype-selective allosteric sites at GPCR-lipid interaction interface

**DOI:** 10.64898/2026.08.06.743397

**Authors:** Jingyi Zhu, Hengde Li, Min Xiao, Lushan Wang, Xukai Jiang

**Affiliations:** State Key Laboratory of Microbial Technology, Shandong University, Qingdao 266237, China; School of Data and Information Sciences, The University of North Carolina at Chapel Hill, Chapel Hill, North Carolina 27514, United States of America; National Glycoengineering Research Center, Shandong University, Qingdao 266237, China

**Keywords:** Multi-agent, GPCR, allosteric modulators, protein-membrane interface

## Abstract

Closely related G protein-coupled receptor (GPCR) subtypes often share highly conserved orthosteric pockets, making subtype-selective ligand development challenging. Here, we developed a five-agent workflow to systematically identify divergent protein-membrane-interface sites across class A GPCRs and exploit them for selective allosteric ligand discovery. By combining dMaSIF-derived surface fingerprints with Ballesteros-Weinstein (BW) position alignment, we compared structurally equivalent membrane-facing regions and identified the three most divergent hotspots for each of 163 receptor pairs. These regions showed substantial spatial overlap with experimentally characterized allosteric sites. Paired target-off-target screening of one million lead-like compounds, followed by detail-mode redocking and multi-seed consistency filtering, yielded 352 receptor-pair-specific candidates corresponding to 344 unique compounds across 104 receptor pairs. These candidates, together with their divergent sites and predicted selectivity profiles, were integrated into a searchable database. Our findings establish a scalable strategy for translating GPCR membrane-interface divergence into precise allosteric sites and testable subtype-selective ligand candidates.

## 1. Introduction

G protein-coupled receptors (GPCRs) constitute the largest family of cell-surface receptors and regulate a broad spectrum of physiological processes, including sensory perception, neurotransmission, metabolism, immune responses, and cardiovascular function^1,2^. Their functional diversity arises not only from the large number of receptor subtypes but also from their ability to populate multiple conformational states and selectively engage heterotrimeric G proteins, β-arrestins, and other intracellular transducers^3,4^. Consequently, GPCRs represent one of the most productive therapeutic target families, with 516 approved drugs acting on 121 GPCRs and accounting for approximately 36% of all approved drugs^2^. However, closely related GPCR subtypes frequently share highly conserved orthosteric binding pockets, making it difficult to achieve subtype selectivity and increasing the risk of unintended modulation of related receptors^5^.

Allosteric sites located outside the endogenous ligand-binding pocket provide an alternative opportunity for selective GPCR modulation^6^. In particular, membrane-facing regions at the receptor-lipid interface can display greater sequence, structural, and physicochemical divergence among related receptor subtypes than their conserved orthosteric pockets, thereby providing a structural basis for subtype-selective ligand recognition^5^. These regions can accommodate endogenous lipids and synthetic modulators that alter receptor conformational equilibria and downstream signalling without necessarily competing directly with orthosteric ligands^7,8^. Moreover, lipophilic ligands can partition into and accumulate within the lipid bilayer, increasing their local concentration around membrane proteins and facilitating lateral access to membrane-exposed binding sites^9^. Membrane-interface sites therefore represent attractive but still underexplored opportunities for expanding GPCR druggability and developing ligands with improved subtype selectivity and potentially prolonged target engagement^10,11^.

Despite the growing interest in targeting receptor-lipid interfaces, GPCR subtype-selective allosteric modulators remain scarce. A recent structure-based survey identified only three approved GPCR allosteric drugs worldwide: avacopan, cinacalcet, and evocalcet^12,13^. Nevertheless, available structural data indicate that membrane-exposed ligand-binding sites are considerably more widespread than suggested by the small number of approved drugs. A recent study curated 72 non-redundant GPCR-ligand structural pairs involving 44 distinct ligand identifiers at protein-lipid bilayer interfaces, highlighting a substantial but largely unexplored chemical space for GPCR modulation^9^. However, receptor or subtype selectivity was not systematically annotated in this dataset. Although several selective membrane-interface ligands have been reported individually, no comprehensive and reproducible estimate of the total number of subtype-selective ligands is currently available. Therefore, systematically identifying subtype-divergent sites at GPCR protein-lipid interfaces and translating these differences into selective allosteric ligand discovery remain major challenges.

Here, we developed a multi-agent computational workflow that integrates GPCR structure preparation, molecular-surface fingerprint sampling, differential hotspot identification, paired virtual screening, and database construction. GPCR structures were standardized using GPCRdb resources, and their molecular surfaces were sampled with dMaSIF to generate local feature representations suitable for cross-receptor comparison^14,15^. Surface features were subsequently aligned according to BW positions, allowing equivalent membrane-facing regions to be compared and the three most divergent regions of each receptor pair to be mapped onto the corresponding three-dimensional structures^16^. The identified hotspots were then subjected to paired virtual screening using Uni-Dock, in which candidate compounds were evaluated against both the intended target and a related off-target receptor to prioritize predicted subtype-selective ligands. Finally, receptor pairs, differential sites, candidate compounds, target and off-target docking scores, and predicted selectivity were integrated into a searchable database, providing a scalable framework for exploiting membrane-interface divergence in GPCR subtype-selective ligand discovery.

## 2. Results

### A multi-agent workflow for the identification of GPCR subtype-selective allosteric modulators

To systematically characterize membrane-interface differences among GPCR subtypes and exploit these differences for selective ligand discovery, we established an automated workflow composed of five sequentially coordinated agents (Fig. 1). The workflow integrates receptor structure standardization, molecular surface fingerprint extraction, differential hotspot identification, paired virtual screening, and database construction, thereby providing a continuous pipeline from GPCR structural inputs to subtype-selective ligand candidates.

**Figure 1.**
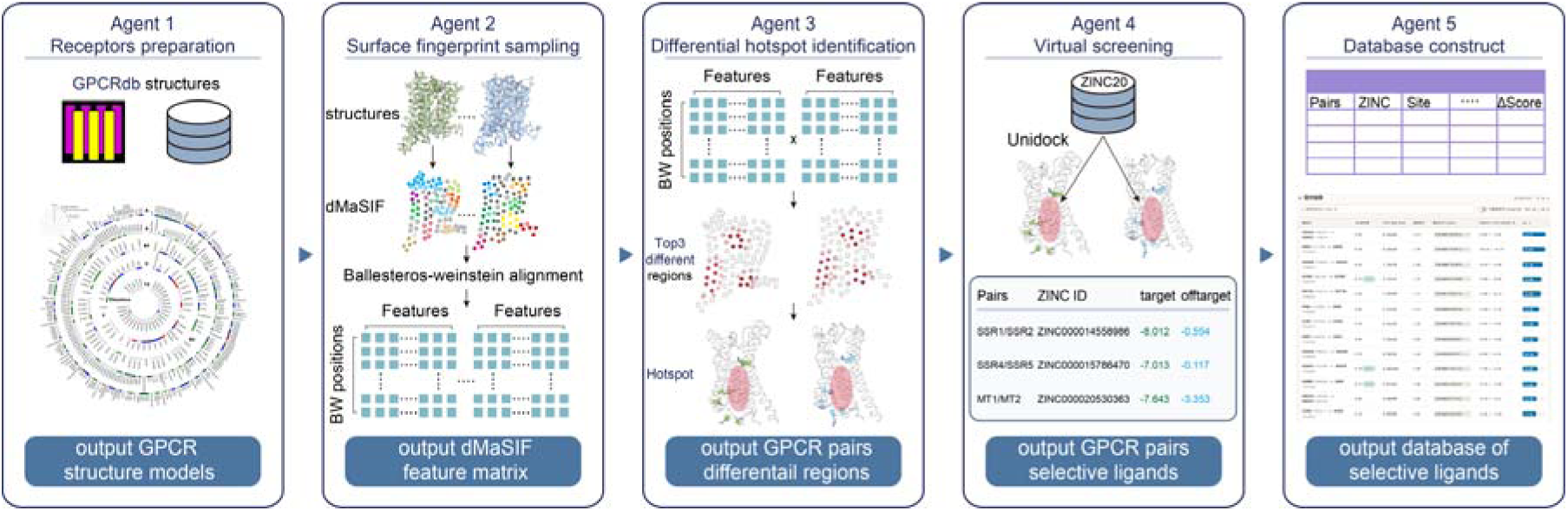
Multi-agent workflow for identifying GPCR membrane-interface differential hotspots and candidate subtype-selective ligands. Agent 1 collects GPCR structures from GPCRdb and performs standardized receptor preparation to generate structural models for downstream analysis. Agent 2 samples receptor molecular surfaces using dMaSIF and encodes local geometric and physicochemical properties as multidimensional surface fingerprints, which are aligned across receptors according to Ballesteros-Weinstein (BW) positions. Agent 3 compares the aligned feature matrices for each receptor pair, identifies the three most divergent membrane-facing regions, and maps them onto the corresponding three-dimensional structures as candidate allosteric hotspots. Agent 4 uses these differential regions as docking sites for paired virtual screening, in which compounds are evaluated against both the target and the corresponding off-target receptor to prioritize candidates with favorable predicted selectivity. Agent 5 integrates receptor pairs, differential sites, compound identities, target and off-target docking scores, and selectivity scores into a structured and searchable database of candidate subtype-selective ligands.

Agent 1 was responsible for receptor preparation. Available GPCR structures were collected from GPCRdb and processed using a standardized procedure to generate receptor models suitable for subsequent surface analysis. This step harmonized structures obtained from different sources and conformational states into a consistent format, thereby establishing a common structural basis for cross-receptor comparison. Agent 2 sampled the molecular surfaces of the prepared GPCR structures using dMaSIF and encoded the local geometric and chemical environment of each surface point as a multidimensional surface fingerprint. The resulting features were subsequently mapped onto corresponding transmembrane positions according to the BW numbering scheme. This procedure generated BW position-indexed surface feature matrices and enabled direct comparison of equivalent membrane-facing regions across different receptors, despite differences in sequence length, local structure, and surface sampling density. Based on these aligned feature matrices, Agent 3 performed position-wise comparisons for each receptor pair and ranked local regions according to their surface-feature differences. The three most divergent regions were selected and mapped back onto the three-dimensional receptor structures. This analysis converted differences in high-dimensional feature space into spatially continuous membrane-facing hotspots with defined structural locations, thereby identifying candidate binding regions that may support subtype-selective ligand recognition. Agent 4 used the identified differential hotspots as docking regions for paired virtual screening against the ZINC20 compound library using Uni-Dock. Each candidate compound was docked independently to both the target receptor and its corresponding off-target receptor. Target and off-target docking scores were recorded simultaneously, and their difference was used to prioritize compounds predicted to bind more favorably to the target receptor while showing weaker binding to the closely related off-target receptor. This paired-screening strategy produced receptor-pair-specific sets of putative selective ligands. Finally, Agent 5 integrated the receptor pair, differential site, compound identity, target and off-target docking scores, and selectivity score into a structured and searchable database. The resulting database established explicit links among receptor pairs, differential membrane-interface sites, candidate compounds, and predicted selectivity, allowing systematic retrieval, comparison, and tracking of screening results.

Collectively, this multi-agent workflow connects GPCR surface-feature comparison with subtype-selective ligand screening in an automated framework. By identifying and exploiting local differences at membrane-facing receptor surfaces, the workflow expands the druggable space beyond conventional orthosteric pockets and provides a systematic computational platform for the discovery of GPCR subtype-selective ligands targeting differential membrane-interface regions.

### Surface fingerprint analysis of Class A non-olfactory GPCRs

Human class A GPCRs comprise 674 receptors, including 388 olfactory receptors (ORs) and 286 non-olfactory receptors. To systematically compare extracellular surface properties across Class A GPCRs, we first curated all non-olfactory Class A receptor structures, representing approximately 42% of the entire Class A GPCR family (Fig. 2a). These receptors span diverse functional groups, with orphan receptors constituting the largest subfamily (79 receptors), followed by chemokine (23), serotonin (11), prostanoid (9), adrenoceptor (8), opsin (7), lysophospholipid (LPA, 6), P2Y (6), dopamine (5), free fatty acid (5), leukotriene (5), lysophospholipid (S1P, 5), and melanocortin receptors (5) (Fig. 2b). To enable quantitative comparison of receptor surfaces, each GPCR structure was processed using dMaSIF, which represents the molecular surface as a point cloud and encodes every surface vertex as a 16-dimensional geometric and physicochemical fingerprint (Fig. 2c). This representation captures local surface characteristics while remaining independent of sequence or structural alignment.

**Figure. 2.**
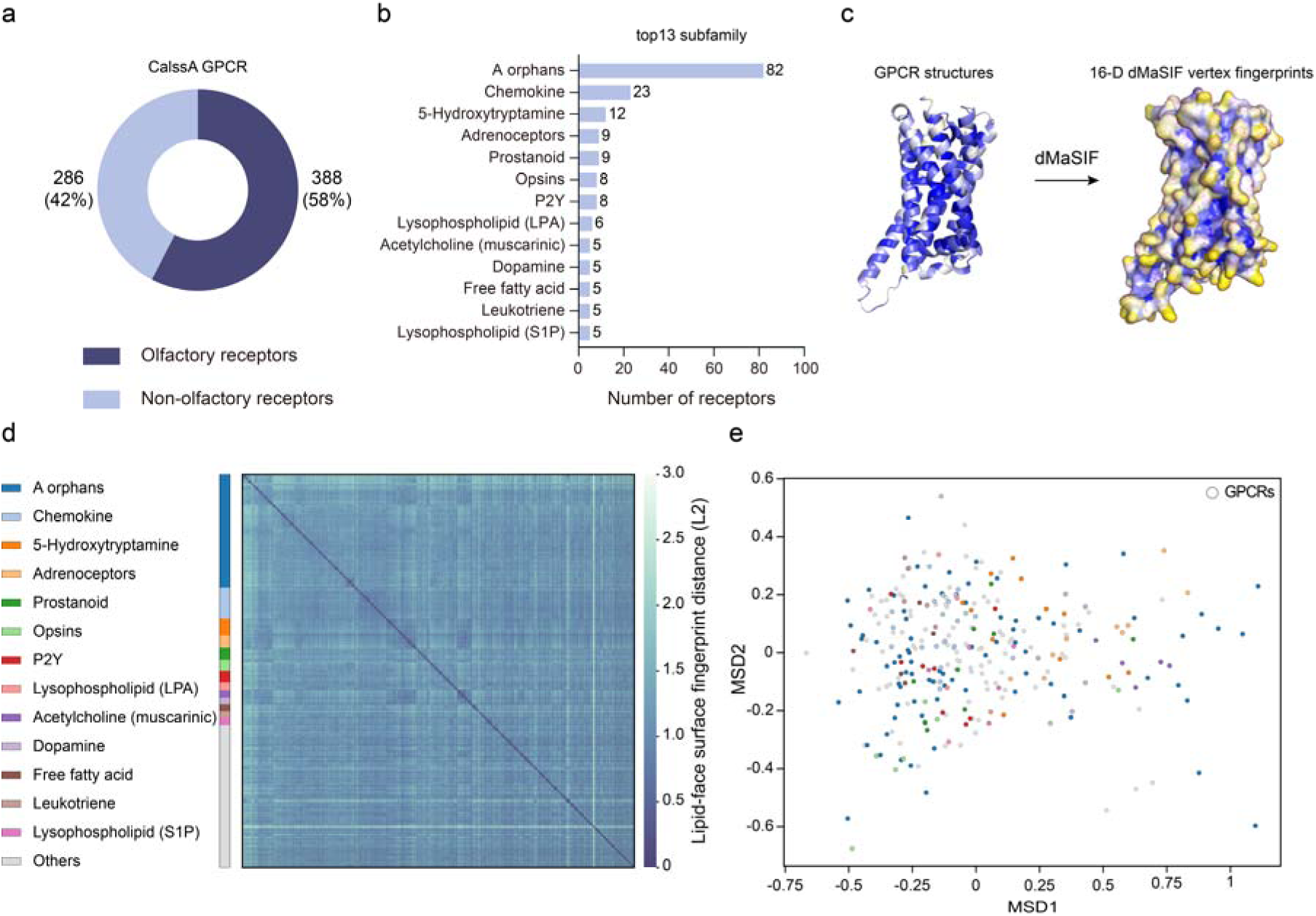
Construction and global analysis of the Class A GPCR surface fingerprint dataset. **a**, Composition of the Class A GPCR family used in this study. Non-olfactory receptors (286 receptors, 42%) were selected for subsequent analyses, whereas olfactory receptors account for 58% of the Class A family. **b**, Distribution of the selected receptors across the 13 largest GPCR subfamilies. The number of receptors in each subfamily is indicated. **c**, Surface fingerprint generation with dMaSIF. GPCR structures were processed using dMaSIF to represent the molecular surface as a point cloud, where each surface vertex was encoded as a 16-dimensional geometric and physicochemical fingerprint. **d**, Pairwise comparison of surface fingerprints among the 286 non-olfactory Class A GPCRs. The heatmap shows pairwise L2 distances between receptor surface fingerprints. Receptors are ordered according to GPCR subfamilies, indicated by the color bar. **e**, Multidimensional scaling (MDS) projection of the pairwise receptor surface embedding distances. Each point represents one GPCR and is colored according to its receptor subfamily.

To enable direct comparison of transmembrane surface fingerprints across receptors, the transmembrane surfaces were first aligned according to the Ballesteros-Weinstein (BW) numbering scheme before pairwise distance calculation. Pairwise distances were then computed between the aligned dMaSIF-derived transmembrane surface fingerprints of all 286 receptors, generating a global similarity (Fig. 2d). The pairwise comparison revealed a high degree of similarity in transmembrane surface fingerprints across Class A GPCRs. Receptors within the same subfamily generally showing higher similarity while substantial overlap was retained across different subfamilies. To further visualize the relationships among receptor transmembrane surface fingerprints, we performed multidimensional scaling (MDS) based on the pairwise distance matrix. In the resulting two-dimensional embedding, each point represents a GPCR, and the distance between two points reflects the dissimilarity of their transmembrane surface fingerprints. Rather than forming discrete clusters, the receptors were continuously distributed in the embedding space (Fig. 2e), providing an intuitive view of the evolutionary relatedness of transmembrane surface properties across Class A GPCRs. This surface fingerprint landscape provides the foundation for subsequent analyses of conserved and subtype-specific allosteric sites.

### Divergent transmembrane surface regions coincide with known allosteric sites

To identify the transmembrane surface regions responsible for receptor-specific differences, we decomposed the global surface fingerprint distance into individual Ballesteros-Weinstein (BW) positions. For each of the 163 GPCR pairs, the top three BW positions exhibiting the largest fingerprint differences were identified (Fig. 3a). These highly divergent positions were distributed throughout the transmembrane bundle but showed clear positional preferences rather than being uniformly distributed. The distribution of the top divergent positions revealed that they were enriched in specific transmembrane helices, particularly TM2, TM3, TM6 and TM7, whereas TM4 and TM5 contributed relatively fewer highly divergent positions (Fig. 3b). These results indicate that receptor-specific membrane interface properties are concentrated within a limited number of transmembrane regions instead of being evenly distributed across the receptor surface.

**Figure 3.**
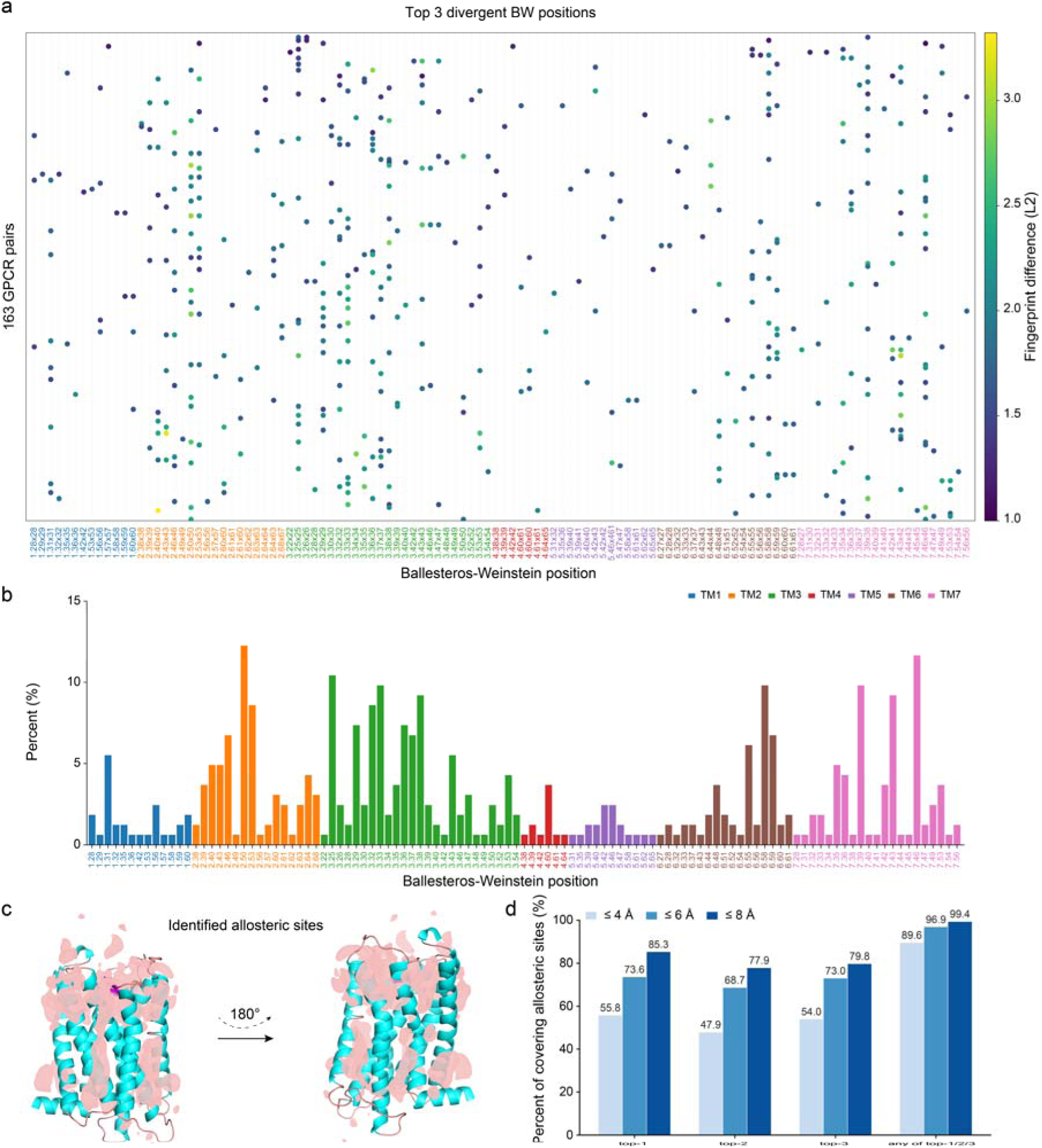
Divergent transmembrane surface regions of class A GPCRs are spatially associated with known allosteric sites. **a**, Distribution of the three most divergent Ballesteros-Weinstein (BW) positions identified for each of the 163 GPCR receptor pairs. Each dot represents one top-ranked divergent position for a receptor pair and is colored according to the L2 distance between the corresponding dMaSIF surface fingerprints. BW positions are grouped and colored by transmembrane helix. **b**, Frequency distribution of the top divergent positions across the transmembrane bundle. Bars indicate the percentage of receptor pairs in which each BW position was identified among the three most divergent positions, with colors denoting TM1-TM7. **c**, Structural mapping of experimentally characterized allosteric sites in class A GPCRs. Allosteric-site regions are shown as pink surfaces on a representative GPCR structure viewed from two opposite membrane-facing orientations. **d**, Spatial coverage of known allosteric sites by the identified divergent regions. Bars show the percentages of experimentally characterized allosteric sites located within 4, 6, or 8 Å of the top-1, top-2, or top-3 divergent region, or within the corresponding radius of any of the three regions. The top-ranked region alone covered 55.8%, 73.6%, and 85.3% of known allosteric sites at 4, 6, and 8 Å, respectively, whereas considering any of the top three regions increased the corresponding coverage to 89.6%, 96.9%, and 99.4%.

To evaluate the functional relevance of these divergent regions, we collected all experimentally reported functional allosteric sites in Class A GPCRs and mapped them onto receptor structures (Fig. 3c). We next quantified the spatial overlap between these divergent regions and experimentally characterized allosteric sites by defining spherical neighborhoods with radii of 4 Å, 6 Å, and 8 Å centered on the top-ranked divergent BW positions (Fig. 3d). The top three divergent regions covered 55.8%, 73.6%, and 85.3% of known allosteric sites within the corresponding radii, respectively. Considering the top three divergent positions further increased the coverage to 89.6%, 96.9%, and 99.4%, respectively. These findings reveal that locally divergent regions of GPCR membrane interfaces are highly enriched at known functional allosteric sites, suggesting that the identified divergent sites represent promising targets for the development of selective allosteric modulators.

### Virtual screening against receptor-pair-specific divergent regions

Based on the receptor membrane-interface divergence analysis described above, we selected the top-ranked divergent region from each of the 163 receptor pairs as the target site for subsequent virtual screening. A library of one million lead-like compounds was first screened using the fast mode of Uni-Dock. Each compound was evaluated at the corresponding divergent sites of both the target and off-target receptors. Predicted selectivity was quantified as the difference in docking score between the target and off-target receptors (ΔΔE=E_target_-E_off-target_), with more negative values indicating a stronger predicted preference for the target receptor. To reduce false-positive predictions arising from insufficient sampling during fast docking, the top 10 candidates were subsequently subjected to symmetric detail-mode redocking against both the target and off-target receptors. Each candidate was evaluated using three independent random seeds, with 10 poses generated for each receptor-seed combination, resulting in a total of 97,800 docking poses. For each seed, the lowest-scoring pose was independently selected for the target and off-target receptors. The detail screening ΔΔE of each candidate was defined as the median of the three seed-specific ΔΔE values. The results showed that more extensive three-seed sampling markedly attenuated the apparent selectivity observed during the fast-screening stage (Fig. 4a). The median ΔΔE shifted from -2.914 kcal/mol in the fast screen to -0.590 kcal/mol after detail-mode redocking, corresponding to a median candidate-wise change of +2.346 kcal/mol. It indicates that detail-mode redocking effectively filtered out potential false-positive candidates arising from insufficient sampling in the fast-mode docking.

**Figure 4.**
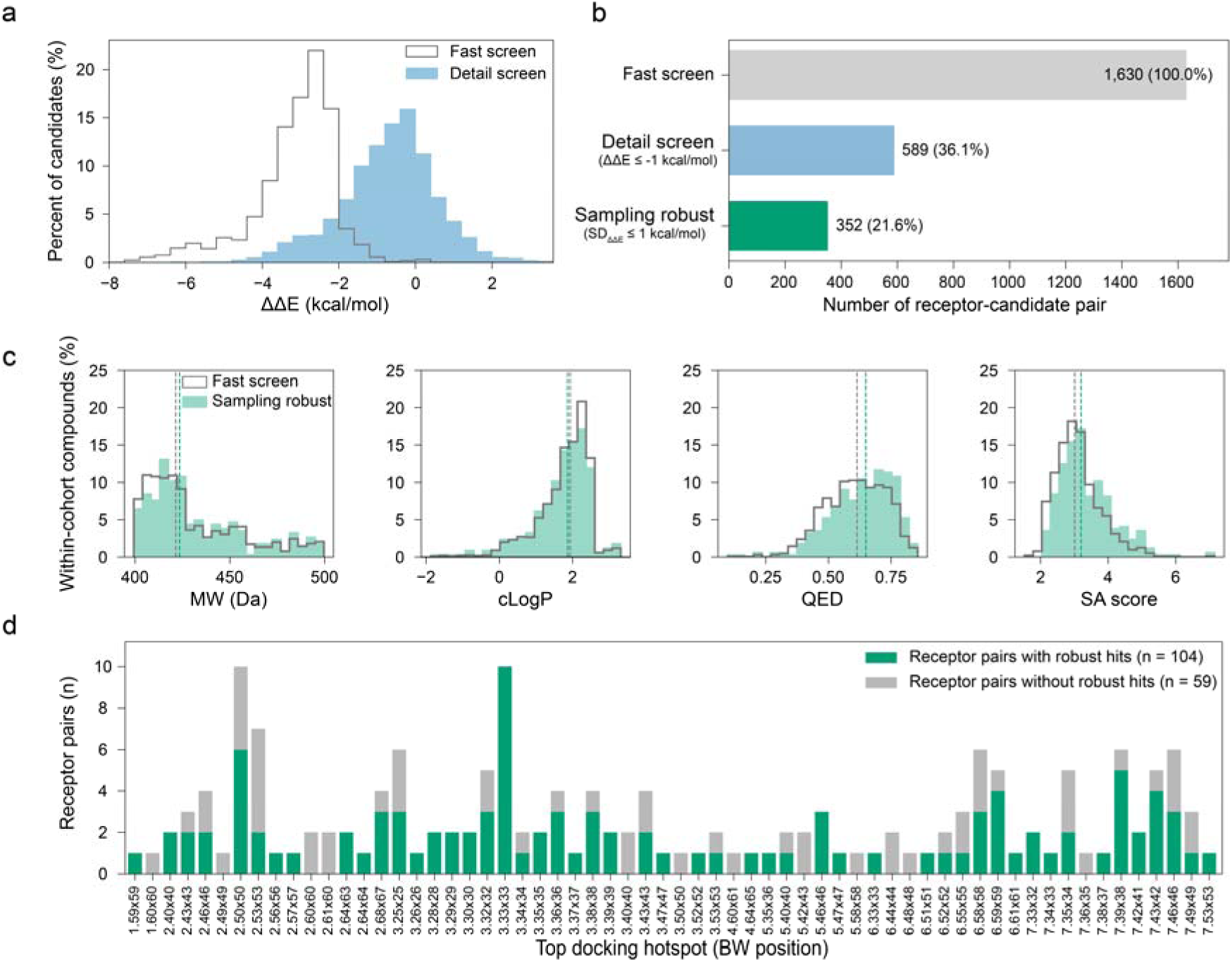
Paired virtual screening and multi-seed filtering of candidate subtype-selective ligands. **a**, Distribution of predicted selectivity scores, defined as ΔΔE=E_target_-E_off-target_, for receptor-candidate pairs obtained from fast-mode screening and subsequent detail-mode redocking. More negative values indicate a stronger predicted preference for the target receptor. **b**, Stepwise filtering of receptor-candidate pairs. The fast screen yielded 1,630 pair-specific candidates. Of these, 589 candidates retained a median detail-mode ΔΔE≤-1 kcal mol^−1^, and 352 candidates further satisfied the multi-seed robustness criterion, defined as an inter-seed standard deviation of ΔΔE<1 kcal mol^−1^. Percentages are relative to the initial candidate set. **c**, Within-cohort distributions of molecular weight (MW), cLogP, quantitative estimate of drug-likeness (QED), and synthetic accessibility (SA) score for compounds from the fast-screen and sampling-robust cohorts. Dashed vertical lines indicate the corresponding cohort medians. The broadly overlapping distributions indicate that multi-seed filtering removed candidates with unstable selectivity predictions without substantially altering the physicochemical-property space. **d**, Distribution of the 163 receptor pairs according to their top-ranked docking-hotspot Ballesteros-Weinstein (BW) positions. Green bars denote receptor pairs retaining at least one sampling-robust candidate (n=104), whereas grey bars denote receptor pairs for which no candidate satisfied all filtering criteria (n=59).

The 10 candidates with the most favorable fast-mode ΔΔE values were retained for each receptor pair, yielding 1,630 pair-specific candidate records (Fig. 4b). Using a detail-mode ΔΔE threshold of ≤ -1 kcal/mol, we identified 589 pair-specific candidates, corresponding to 36.1% of the initial candidate pool. These candidates were further evaluated for sampling robustness, which was defined as an inter-seed standard deviation (SD) of < 1 kcal/mol across the three independent docking runs. This additional criterion retained 352 candidates, representing 59.8% of the 589 selective candidates and 21.6% of the initial candidate pool. Application of the multi-seed consistency criterion further eliminated potential false-positive candidates arising from sampling variability. The 352 pair-specific candidates corresponded to 344 unique ZINC compounds. Their molecular weight (MW), cLogP, quantitative estimate of drug-likeness (QED), and synthetic accessibility (SA) score distributions largely overlapped with those of the 1,522 unique compounds obtained from the fast screen (Fig. 4c). The median values for the sampling-robust and fast-screen cohorts were 423.42 and 421.50 Da for molecular weight, 1.86 and 1.93 for cLogP, 0.65 and 0.61 for QED, and 3.20 and 3.01 for the synthetic accessibility score, respectively. These results indicate that multi-seed validation primarily removed candidates with unstable selectivity predictions without substantially shifting the overall physicochemical-property space. At the receptor-pair level, 104 of the 163 receptor pairs retained at least one sampling-robust candidate, whereas the remaining 59 pairs contained no candidates satisfying all selection criteria (Fig. 4d). The two groups of receptor pairs were distributed across a broad range of top docking-hotspot BW positions. The retained candidates originated from divergent sites distributed across multiple BW positions, with TM3-associated sites yielding the largest number of sampling-robust candidates. These compounds were subsequently used as starting points for further lead optimization.

## 3. Discussion

Despite the growing recognition of the protein-lipid interface as a promising source of GPCR allosteric sites, the development of selective ligands targeting these regions remains substantially underexplored^8–10^. Compared with conventional orthosteric pockets, membrane-facing sites are often shallow, conformationally dynamic, and partially occupied by lipids, making them difficult to identify and exploit using established structure-based drug-discovery strategies^9,17,18^. Moreover, although individual membrane-interface modulators have demonstrated that these sites can support receptor or subtype selectivity, a systematic framework for comparing equivalent membrane-facing regions across the GPCR superfamily and translating their local differences into selective ligand discovery has been lacking. In this study, we developed a multi-agent workflow that integrates molecular-surface fingerprinting, generic-position-guided cross-receptor comparison, differential hotspot mapping, and paired target-off-target virtual screening. Application of this framework to 286 non-olfactory class A GPCRs and 163 receptor pairs revealed broadly distributed membrane-interface differences and generated receptor-pair-specific candidate ligands, supporting the GPCR membrane interface as an underutilized but potentially generalizable space for selective allosteric ligand discovery.

Through systematic comparison of membrane-facing molecular surfaces, we identified subtype-divergent hotspots across class A GPCRs that are not readily captured by conventional sequence alignment or pocket-detection approaches. Previous large-scale GPCR analyses have primarily focused on sequence conservation, orthosteric ligand recognition, receptor activation, or intracellular transducer coupling, whereas the local geometric and physicochemical diversity of the receptor-lipid interface has received less systematic attention. By encoding receptor surfaces with dMaSIF-derived fingerprints and aligning structurally equivalent regions according to Ballesteros-Weinstein positions, we directly compared corresponding membrane-facing environments across 286 non-olfactory class A GPCRs. Analysis of 163 receptor pairs revealed three highly divergent regions for each pair, which were subsequently mapped onto the receptor structures as spatially defined hotspots. Notably, these regions showed substantial spatial overlap with previously reported functional allosteric sites, suggesting that local surface divergence is frequently associated with chemically accessible regulatory regions. Thus, rather than defining potential binding sites as broad transmembrane segments, our analysis provides precise three-dimensional locations that can be directly used for subtype-selective allosteric ligand screening and structure-guided optimization.

Building on these structurally defined hotspots, we performed paired virtual screening to translate receptor-surface divergence into predicted chemical selectivity. Conventional virtual screening typically evaluates compounds against a single receptor, and therefore prioritizes predicted target affinity without directly assessing recognition of closely related off-target subtypes^5,19,20^. In our workflow, each compound was docked independently to both members of a receptor pair, allowing target-off-target differences to be evaluated under the same screening conditions. The initial screen identified 1,630 receptor-pair-specific candidates, of which 589 retained a median ΔΔE of ≤ -1 kcal/mol after detail-mode redocking. Application of multi-seed consistency criteria further identified sampling-robust candidates for 104 of the 163 receptor pairs, indicating that membrane-interface differences can be converted into testable ligand-selectivity hypotheses for a substantial proportion of class A GPCR pairs. We subsequently organized the receptor pairs, divergent hotspots, candidate structures, docking poses, target and off-target scores, and selection outcomes into a searchable database. This organization preserves the relational nature of selectivity, because the predicted preference of a compound is defined with respect to a particular target-off-target pair rather than to an isolated receptor. The database therefore provides a resource for retrieving candidate selective allosteric ligands, comparing chemical solutions across divergent membrane sites, and prioritizing compounds for subsequent simulation, experimental validation, and lead optimization.

Several limitations should be considered when interpreting these results. The current analysis is primarily based on static receptor structures and docking-derived selectivity estimates, which may not fully capture receptor conformational dynamics, lipid competition, or ligand behaviour within the membrane. In addition, the identified compounds remain computational candidates and require validation through molecular dynamics simulations, binding assays, mutagenesis, and functional experiments. Incorporating receptor conformational ensembles and experimental feedback will further refine hotspot identification and compound prioritization.

Overall, our framework provides a scalable strategy for translating subtype-divergent GPCR membrane interfaces into structurally defined candidate allosteric sites and experimentally testable hypotheses of ligand selectivity. By linking cross-receptor surface comparison with paired target–off-target screening, the workflow moves beyond the identification of local structural differences and directly connects these differences to potential chemical recognition. The resulting hotspot and ligand database further enables systematic retrieval and comparison of receptor-pair-specific candidates, providing a foundation for subsequent molecular simulation, experimental validation, and lead optimization. These findings support the GPCR protein-lipid interface as a broadly distributed yet underexploited chemical space for the development of selective allosteric modulators beyond conserved orthosteric pockets.

## 4. Materials and Methods

### Workflow architecture and reproducibility controls

We developed a six-agent computational workflow for identifying subtype-selective ligands targeting membrane-exposed surfaces of class A GPCRs. A typed shared state connected the agents in the following order: receptor curation (Agent 1), structural representation (Agent 2), differential surface analysis (Agent 3), molecular screening and design (Agent 4), evidence-based triage (Agent 5). Human-review checkpoints were placed after Agents 1, 3 and 5.

The orchestration layer was configured with Claude-family large language models. Claude Sonnet 4.6 was assigned to Agents 1, 2, 4 and 6, whereas Claude Opus 4.7 was assigned to Agents 3 and 5. Temperatures ranged from 0 to 0.3. Language models were restricted to evidence synthesis, task planning and structured interpretation. Receptor pairing, surface-distance calculation, docking, pose analysis, physicochemical filtering, ADMET prediction, MM/GBSA calculation and Pareto selection were performed using deterministic programs. Language-model output could not directly override numerical eligibility gates. Each production stage generated a machine-readable manifest containing software versions, configuration parameters, input identities, file hashes and output counts. SHA-256 hashes were used to verify scripts, model checkpoints, input bundles and result files. Downstream execution was halted after any non-zero exit, missing output, non-finite value, cache inconsistency or contract mismatch.

### Agent 1: receptor-universe definition

Agent 1 constructed the receptor universe from human class A GPCRs annotated in GPCRdb^21^. The resulting production universe comprised 286 non-olfactory human class A GPCRs with AlphaFold structural models^22,23^. GPCRdb generic residue numbers were used to provide a common transmembrane coordinate system. This avoided direct comparison of raw residue indices between receptors of different lengths. Agent 1 also maintained receptor identities, UniProt accessions, GPCRdb annotations and structural-source provenance for all downstream analyses.

### Agent 2: structural standardization and surface representation

Agent 2 retrieved and standardized the AlphaFold model for each receptor. Molecular surfaces were encoded using the pretrained dMaSIF model dMaSIF_search_3layer_12A_16dim^14^. For every surface vertex, the 16-dimensional learned descriptor produced by the final feature layer was retained. The transmembrane bundle axis was estimated from the receptor structure. Surface vertices were classified as lipid-facing using the cosine between the vertex normal and the radial vector from the bundle axis. Vertices with a radial-normal cosine greater than 0.30 were considered lipid-facing. For each GPCRdb generic position, a local surface patch was defined using vertices within 6 Å of the corresponding Cα atom. A generic position was retained only when both receptors being compared contained at least three lipid-facing surface vertices within the patch. The descriptor for each retained position was calculated as the mean 16-dimensional dMaSIF vector across its lipid-facing vertices. This production analysis used the dMaSIF-based membrane-surface representation. A conventional cavity-detection module was present in the general Agent 2 architecture but was not used to define the receptor pairs reported here.

### Agent 3: receptor-pair and differential-hotspot identification

Agent 3 compared the lipid-facing surface descriptors of all unordered receptor pairs. For a shared generic position k, the local difference was defined as the Euclidean distance between the two 16-dimensional descriptor vectors:

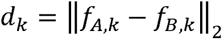

The receptor-level surface distance was calculated as the mean of d_k_ across all supported shared positions. Receptor pairs with a mean surface distance below 0.8 were retained, producing 163 unordered receptor pairs.

Within each pair, shared generic positions were ranked by decreasing local descriptor difference. The highest-ranking position was used as the primary docking hotspot. The docking-box centre was placed 4 Å radially outward from the corresponding Cα atom relative to the transmembrane-bundle axis. The three highest-ranking positions were additionally retained for interpretation and structural reporting. The dMaSIF distance was treated as an exploratory membrane-surface descriptor rather than an experimentally validated measure of receptor selectivity.

### Agent 4: bidirectional small-molecule screening

Agent 4 evaluated each of the 163 receptor pairs in both directions, giving 326 directed target/off-target tasks. For each directed task, 1,000,000 purchasable three-dimensional compounds were sampled from the ZINC20 in-stock collection^24^. Initial docking was performed with Uni-Dock in fast mode using the Vina scoring function^25^. Target and off-target structures were screened independently using the same compounds and a 22 × 22 × 22 Å docking box. Directional selectivity was defined as

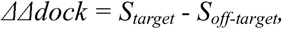

where a more negative value indicated stronger predicted binding to the designated target. The ten highest-ranked compounds from each directed task were retained. Additional compounds from ranks 11-30, 31-100 and, where required, 101-200 were evaluated to improve directional coverage.

Detailed docking was then performed with Uni-Dock in detail mode. Each ligand was docked with random seeds 17,171, 29,292 and 43,434, generating ten poses per seed. A compound passed the formal directional docking gate only when its seed-matched ΔΔdock was no greater than -1.0 kcal mol^-1^ for all three seeds and the population standard deviation across seeds did not exceed 1.0 kcal mol^-1^. Fast-mode docking was used only for enrichment and could not independently qualify a compound. Pose convergence was evaluated in the receptor coordinate frame. The best poses from the three seeds were required to have a maximum centroid separation of 2.5 Å, a median 2 Å voxel-occupancy Jaccard index of at least 0.35, and a median receptor-contact Jaccard index of at least 0.50. The largest cluster among poses within 2 kcal mol^-1^ of the global minimum had to contain at least 50% of the near-best poses. The standard deviation of the three seed-best docking scores also had to be no greater than 1.0 kcal mol^-1^.

### Agent 5: deterministic evidence triage

Agent 5 integrated directional docking robustness, pose convergence, chemical identity and provenance evidence. It did not use a weighted composite score. Compounds were first required to pass the three-seed directional docking gate. A formal seed was then selected for each eligible direction using a frozen lexicographic rule: lowest worst-seed ΔΔdock, lowest seed standard deviation, lowest median detailed-docking ΔΔdock, best fast-screen rank and, finally, lexical ZINC identifier. The bidirectional analysis produced 304 formal directional seeds. Of these, 244 passed the complete pose and chemical-identity audit and entered Agent 6. Fifty-two were retained as pose-consensus holds, and eight were retained as chemical-mapping holds. Held compounds were preserved in the audit ledger but were not treated as eligible lead-optimisation seeds.

### Physicochemical and ADMET evaluation

Physicochemical descriptors were calculated using RDKit. The hard property limits were molecular weight 200-600 Da, cLogP -0.5 to 6.0, topological polar surface area 15-140 Å², no more than 12 rotatable bonds, QED of at least 0.35 and synthetic-accessibility score no greater than 5.0. ADMET properties were predicted using ADMET-AI version 1.4.0 with ten frozen model files and 98 endpoints^26^. The principal safety endpoints were Ames mutagenicity, clinical toxicity, drug-induced liver injury, hERG inhibition and inhibition of CYP1A2, CYP2C19, CYP2C9, CYP2D6 and CYP3A4. Exposure-related endpoints included predicted oral bioavailability, human intestinal absorption and PAMPA permeability. Aqueous solubility was assessed using the AqSolDB endpoint. A candidate was seed-relative eligible only when its maximum predicted liability across the nine safety endpoints did not exceed that of its matched seed. It also had to improve strictly in at least one of three attributes: QED, synthetic-accessibility score or distance from the preferred cLogP interval of 1-5.

### Paired MM/GBSA rescoring

Candidate and matched-seed complexes were subjected to the same frozen MM/GBSA protocol using AmberTools 24.8^27^. Protein and ligand parameters were assigned using ff14SB and GAFF2, respectively. AM1-BCC ligand charges were generated once per compound and shared across receptor roles and pose clusters. The mbondi2 radii set was used. Complexes were minimized with sander in two stages. The protein backbone atoms C, Cα, N and O were restrained. Stage 1 comprised 500 cycles, including 100 steepest-descent cycles, with a restraint of 10 kcal mol^-1^ Å^-2^. Stage 2 comprised 1,500 cycles, including 300 steepest-descent cycles, with a restraint of 2 kcal mol^-1^ Å^-2^.

Binding energies were estimated from one minimised snapshot using MMPBSA.py, the GBOBC model (igb=5), a salt concentration of 0.150 M and a non-bonded cut-off of 999 Å. Entropic contributions were not calculated. The standard AM1 calculation used grms_tol=0.0005, scfconv=1 × 10^-10^ and 700 DIIS attempts. An iteration limit of 5,000 was used only after an exact initial AM1 failure caused by 1,000-step SCF non-convergence.

MM/GBSA selectivity was defined as

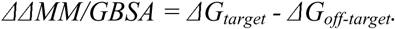

Candidate improvement was assessed relative to the corresponding seed processed with the same runner and receptor contract. Negative candidate-minus-seed values indicated improved target selectivity. Both median-pose and worst-pose improvements were required to be negative for terminal paired advancement.

### Pareto selection and synthesis assessment

Eligible candidates were evaluated in three independent Pareto domains. The docking domain minimized detailed ΔΔdock, seed standard deviation and median target docking score. The drug-like domain maximized QED while minimizing synthetic-accessibility score and molecular weight. The ADMET domain minimized maximum safety liability while maximizing the exposure floor and predicted solubility.

A compound entered the shortlist only when it was seed-relative eligible and Pareto-optimal in at least two of the three domains. ECFP4 similarity to the seed was reported as an interpretive descriptor but was not used as a terminal ranking score. No weighted total score was used.

For shortlisted compounds, retrosynthetic accessibility was assessed using AiZynthFinder 4.4.1 with the official USPTO expansion policy, reaction templates and quick-filter model. Searches were restricted to commercially available Mcule building blocks. Initial searches used a maximum depth of three transformations and 500 iterations. Unsolved compounds could undergo frozen, non-overwriting searches using up to 5,000 iterations and, subsequently, a maximum depth of five transformations. Selected unresolved cases were cross-checked using Syntheseus with LocalRetro. Computational routes were treated as prioritisation evidence and not as experimental demonstrations of synthetic feasibility.

## Acknowledgments

This work was supported by the National Key Research and Development Program (2023YFC3403502), National Natural Science Foundation of China (32301041, 32571437), TaiShan Scholar (NO.tsqn202507080), SKLMT Frontier and Challenges Project (SKLMTFCP-2023-01) and SKLDRS Open Project (2025SKLDRS0323), Intramural Joint Program Fund of State Key Laboratory of Microbial Technology (Project NO. SKLMTIJP-2025-03).

